# Non-invasive vagus nerve stimulation modulates Pavlovian bias in a state-dependent manner

**DOI:** 10.64898/2025.12.14.693973

**Authors:** Luisa Kaluza, Anne Kühnel, Nora Gerth, Yul Wegner, Corinna Schulz, Nils B. Kroemer

**Author notes:** Corresponding author: Prof. Dr. Nils B. Kroemer, Venusberg Campus 1, 53127 Bonn, Germany.

## Abstract

The vagus nerve transmits vital signals between organ systems of the body and the brain. Despite growing interest in non-invasive transcutaneous vagus nerve stimulation (tVNS) for treatments of various disorders, it is not known whether its modulatory effects on behavior are dependent on bodily states. Here, we used a single-blind randomized crossover design with two within-participant factors, stimulation (right tVNS vs. sham) and metabolic state (water vs. caloric load), to investigate 54 healthy participants (210 sessions). To evaluate state-dependent changes in reinforcement learning, we used a valenced go/no-go task. Performance on this task can be used to estimate a Pavlovian choice bias, which captures improved performance for valence-congruent actions (e.g., go to win rewards) versus valence-incongruent actions (e.g., no-go to win rewards) and has been associated with dopaminergic neurotransmission. In line with theorized state-dependency of tVNS-induced changes, we observed that tVNS decreased Pavlovian response biases (Go × Win) after the caloric load (Stimulation × Load: *p* =.038). Taking into account individual changes in hunger but not satiety ratings best captured behavioral modulation, indicating that this modulatory effect could be driven by motivational effects related to metabolic interoception (sensed “need”). We conclude that right tVNS decreases Pavlovian response biases if the body is in a postprandial, but not a fasting state. Since fasting and Pavlovian biases have been previously associated with elevated dopamine neurotransmission, our results call for additional research on state-dependent motivational effects of tVNS to optimize interventions.

## Introduction

To ensure survival, organisms must learn to adapt their behavior based on current homeostatic needs, for example, by pursuing options that maximize energy intake during starvation. Vagal afferent signaling has often been conceptualized as primarily transmitting negative homeostatic feedback to the brain (Bai et al., 2019; de Lartigue, 2016; Young et al., 1974; Yox et al., 1991). However, recent evidence suggests that the role of the vagus nerve extends beyond negative feedback. In addition to suppressing food intake, vagal inputs to the nucleus tractus solitarii (NTS), the primary target of vagal afferents in the brain, potentially affect neural populations that promote food-seeking behavior (Chen et al., 2020; Shen et al., 2025). Additionally, rodent studies provide evidence of state-dependent vagal signaling by demonstrating vagally mediated stomach–brain coupling according to metabolic state (Cao et al., 2022). Thus, recent evidence suggests that the vagus nerve fine-tunes allostatic behavior along with the body’s state (Teckentrup and Kroemer, 2024), activating different neural and signaling pathways as needed to support appropriate behavior. Notably, the vagus nerve links food intake with dopaminergic reinforcing signals (Buchanan et al., 2022; Ferreira et al., 2012; Kaelberer et al., 2018; Ren et al., 2010; Weber et al., 2025). Recent evidence has further linked vagal signaling with dopaminergic signaling more broadly, indicating an influence that extends beyond natural rewards (Onimus et al., 2025). Collectively, these findings raise the question of whether internal bodily states need to be considered in targeted interventions of the vagus nerve aimed at modulating reward-related behavior.

Transcutaneous auricular vagus nerve stimulation (tVNS) modulates vagal activity via the auricular branch of the vagus nerve (Fallgatter et al., 2003), leading to robust activation of the NTS (Frangos et al., 2015; Teckentrup et al., 2021). Downstream from the NTS, VNS has been shown to induce widespread neuromodulatory effects on various neurotransmitters, such as noradrenaline (Roosevelt et al., 2006), dopamine (Brougher et al., 2021; Han et al., 2018; Onimus et al., 2025), and acetylcholine (Bowles et al., 2022; Hulsey et al., 2016; Mridha et al., 2021). Monoaminergic signaling in the brain is a key mechanism regulating reinforcement learning (Beeler, 2012; Pessiglione et al., 2006; Schultz et al., 1997; Steinberg et al., 2013) and approach (vs. avoidance) behavior (Beierholm et al., 2013; Hamid et al., 2016; Niv et al., 2007; Rigoli et al., 2016; Wise, 2004). In addition, dopamine signaling has been linked to Pavlovian biases (Eisinger et al., 2020; Guitart-Masip et al., 2014; Scholz et al., 2022), which reflect better performance on valence-congruent (e.g., go to win) versus incongruent (e.g., no-go to win) actions (Guitart-Masip et al., 2012b; Hershberger, 1986). In line with the neuromodulatory effects, VNS has been shown to affect motivation and learning in rodents (Bowles et al., 2022; Brougher et al., 2021; Han et al., 2018; Sahasrabudhe et al., 2023) and humans (Çakır et al., 2025; Ferstl et al., 2024; Neuser et al., 2020; Weber et al., 2021). However, other studies reported decreased learning rates or no effect on learning performance or effort invigoration using tVNS in humans (D’Agostini et al., 2021; Kühnel et al., 2020; Lucchi et al., 2024; Thanarajah et al., 2025). Such acute effects of VNS are likely dependent on internal and external states because VNS elicits a fear-state dependent effect on extinction performance (Klein et al., 2021) and improvements in motor learning depend on the task state in rodents (Bowles et al., 2022). This is substantiated by findings in humans demonstrating influences of metabolic state and interoception on tVNS-induced effects (Kozorosky et al., 2022; Salaris and Azevedo, 2024). Since right VNS activates dopaminergic projections more robustly (Brougher et al., 2021; Han et al., 2018), while human studies on reinforcement learning predominantly used left tVNS without considering metabolic states (Çakır et al., 2025; D’Agostini et al., 2021; Weber et al., 2021), these open questions hamper the optimization of tVNS-based treatments.

To address these gaps, we investigated whether metabolic state modulates the effects of right tVNS on reinforcement learning using a randomized crossover design. To this end, we manipulated metabolic state within participants with a load manipulation (milkshake vs. water) prior to right tVNS (vs. sham). Healthy participants (*N* = 54) then completed a valenced go/no-go task (Guitart-Masip et al., 2014, 2012b; Mkrtchian et al., 2017). In line with previous work (Kühnel et al., 2020), we found a tVNS-induced reduction of the learning rate, but only in a fasting state. Crucially, in line with the expected motivational effects, we observed tVNS-induced changes of the Pavlovian bias depending on the caloric load before the task, leading to increases after water and decreases after milkshake. This modulatory effect was best captured by individual changes in hunger but not satiety ratings, suggesting that the interoceptive sensing of the caloric load may drive “need” state-dependent effects of right tVNS.

## Methods

### Participants

Initially, we enrolled 74 physically and mentally healthy participants, as determined by phone screenings. For the reported analyses, we excluded 20 participants (*n* = 10 dropped out before the first session, *n* = 5 dropped out before completion of two sessions, *n* = 2 before completion of all four sessions, and *n* = 3 had go responses in <5% of all trials). In addition, we had to exclude 6 sessions due to other reasons (2 sessions were affected by equipment error leading to missing go responses, 2 sessions were affected by errors in the stimulation protocol, 2 sessions were affected because participants initially used the wrong device to respond). Consequently, we analyzed data from 54 participants (30 women, *M_age_* = 35.3 ± 13.7 years, *M_BMI_* = 28.0 ± 5.94 kg/m^2^) and 210 individual sessions. The study was approved by the local ethics committee according to the Declaration of Helsinki and each participant gave written informed consent prior to participation.

### Experimental procedure

The study used a single-blind, randomized crossover design with two within-participant factors: right tVNS (vs. sham) and metabolic state (water load vs. caloric load) leading to four sessions for each participant (Fig. 1A). Sessions took place after an overnight fast (*M* = 13.8 ± 1.83 h) between ∼08:00 am and ∼2:30 pm, with about one week in between sessions (*M* = 7.22 ± 6.92, range 3–51 d). Then, each session started with anthropometric measurements such as weight, height, hip and waist circumference, as well as resting pulse rate. This was followed by a first set of state questions (e.g., hunger and satiety) that participants completed on a visual analog scale (VAS T1; Ferstl et al., 2022). Next, participants received either a high-calorie milkshake (∼500 kcal) or an equal volume of water, which had to be consumed within ∼5 min. Throughout the sessions, participants could drink as much water as they liked. After ∼15 min, participants completed the state questions a second time (VAS T2). Afterward, tVNS or sham was applied according to the randomization list (generated in advance; participants were assigned after enrolment). After calibration of the stimulation strength, the stimulation (30s ON, 30s OFF) continued throughout all task blocks. Within this block, participants completed a food cue reactivity task (∼20 min; Müller et al., 2022), another set of state questions (VAS T3), and an effort allocation task (∼40 min; Ferstl et al., 2024; Neuser et al., 2020) before the valenced go/no-go learning task (∼20 min; Guitart-Masip et al., 2014, 2012b; Kühnel et al., 2020; Mkrtchian et al., 2017). After the task, participants completed the state questions again (VAS T4) and then received rewards according to their task performance, including their breakfast according to the energy earned in the effort task. At the end, participants completed the state questions again (VAS T5). After the last session, participants received compensation (either as 80 € or partial course credit) and each session followed the same standardized procedure.

**Figure 1:**
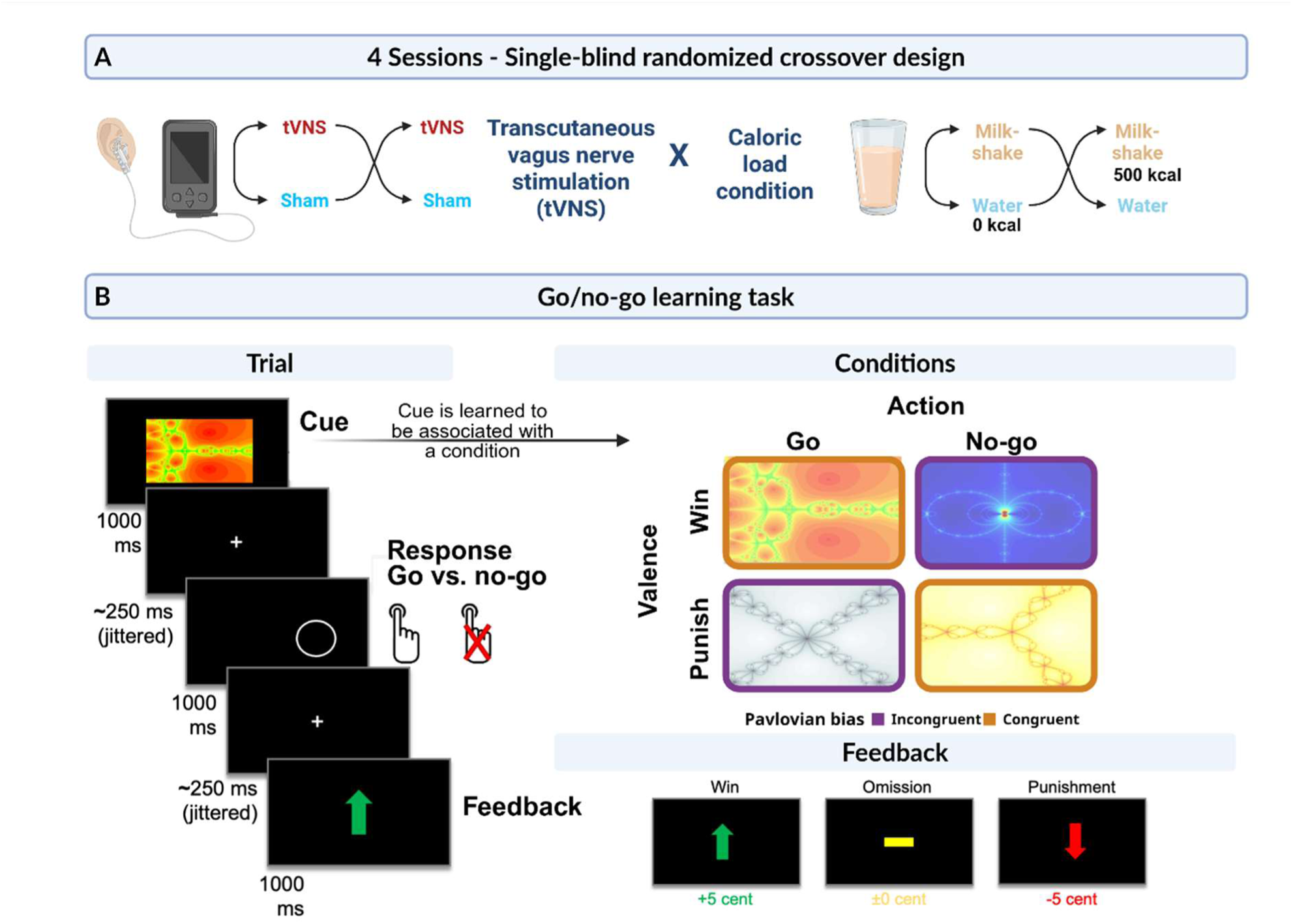
Study and task design. A: The study used a single-blind, randomized crossover design with two within-subject factors: stimulation (right tVNS vs. sham) and metabolic state (water load vs. caloric load), resulting in four sessions per participant. B: In the task, participants learn state-action contingencies by trial and error. In each trial, participants see a fractal cue randomized to one of the four possible combinations (Go vs. No-go × Win vs. Punishment). Participants then respond by either pressing a button (go) to complete a target detection task or withholding their response (no-go), which is then followed by the outcome presentation (win, punishment, or omission).

### Transcutaneous vagus nerve stimulation device

We used the Cerbomed NEMOS device (Cerbomed GmbH, Erlangen), which delivers a 250 µs pulse width, 25 Hz stimulation in a biphasic 30 s ON, 30 s OFF duty cycle, in line with previous studies (Ferstl et al., 2024; Teckentrup et al., 2021). For tVNS, the electrode was placed on the right ear at the cymba conchae targeting the auricular branch of the vagus nerve (Frangos et al., 2015). For sham stimulation, we used the earlobe, which is not innervated by the vagus nerve (Farmer et al., 2021; Peuker and Filler, 2002). To ensure adequate skin contact, we rubbed the skin with alcohol and placed medical tape to secure the electrodes in place.

Stimulation strength was set individually and separately per session using a pain VAS rating scale (“How intensely do you feel pain induced by the stimulation?” ranging from 0 (“no sensation”) to 10 (“strongest sensation imaginable”)). Stimulation was initiated at an amplitude of 0.1 mA and increased by the experimenter in 0.1–0.2 mA steps. Participants rated the sensation after every increment until ratings settled around 5, corresponding to “mild pricking”. The stimulation strength was *M_tVNS_* = 1.77 ± 0.63 mA and *M_sham_* = 2.10 ± 0.55 mA.

### Valenced go/no-go learning task

To investigate whether tVNS effects on reinforcement learning depend on metabolic state, we used a previously established orthogonalized go/no-go learning task (Guitart-Masip et al., 2014, 2012b; Kühnel et al., 2020; Mkrtchian et al., 2017; Fig. 1B). In the task, participants learn state-action contingencies by trial and error from receiving rewards (5 cents), punishments (-5 cents), or neutral outcomes (0 cents). Each trial comprises three stages. In the first stage of each trial, participants see a fractal cue (state) randomized to one of the four possible combinations (Go vs. No-go × Win vs. Punishment). Second, participants had to either respond by pressing a button (go) to subsequently complete a target detection task or withhold their response (no-go). In the third stage, the outcome was presented to the participant, which was either a win, a punishment, or an omission. The assigned outcomes were presented probabilistically. Accordingly, there was an 80% chance to either win or avoid losses after correct state–action sequences, and a 20% chance to win or avoid losses after incorrect sequences. In total, the task included 240 trials, with 60 trials for each condition, and took about 20 min to complete. The probabilistic nature of the task and that either go or no-go responses could be correct for a given fractal were explained to the participants prior to task performance.

## Data analysis

### Model-agnostic statistical analysis

We defined a full linear mixed-effects model to estimate the effects of tVNS and caloric load on choice accuracy. To model task conditions, we included the regressors go (centered), win (centered), their interaction (Go x Win), a trial regressor to capture learning (ln-transformed and centered), and session (ln-transformed and centered) to capture improvements over sessions. To assess tVNS and metabolic-state effects, the model included regressors for the stimulation condition (effect centered:-0.5 = sham, 0.5 = tVNS), the caloric load condition (effect centered:-0.5 = water, 0.5 = milkshake), their interaction, and interactions with the task regressors. The model included random intercepts and random slopes for the regressors go, win, stimulation, load, their interaction, and the trial regressor per ID (for model equations, see SI1).

We tested an additional model including three centered participant-level predictors (sex, age, and BMI), which did not change the reported results (Table SI2). In addition, we also tested whether tVNS-induced effects were partly dependent on BMI as previously reported (Kühnel et al., 2020), by extending the interaction of Go × Win with Stimulation × Load in an additional model (Table SI3).

Since identical energy intake may elicit variable metabolic responses across individuals (Theodorakis et al., 2024), we also used individual changes in hunger and satiety ratings as a proxy for perceived changes in metabolic state after the load (Kaduk et al., 2025; Lemmens et al., 2011; Schultes et al., 2003). In line with the concept of distinct vagal pathways to promote or suppress food seeking (Chen et al., 2020; Shen et al., 2025), we estimated two models including individual load-induced changes in hunger (ΔHunger = (Hunger T2 – Hunger T1) / 100) and satiety (ΔSatiety = (Satiety T2 – Satiety T1) / 100) instead of the load condition, respectively. To assess the effects of tVNS on Pavlovian bias (Go × Win) when taking hunger or satiety into account, we used the random slopes of the Go × Win regressor as the dependent variable and included the centered regressors stimulation and hunger or satiety, as well as their respective interactions. The model included random intercepts and random slopes for stimulation and hunger/satiety per ID.

### Reinforcement learning model

To identify whether a specific facet of learning was affected by right tVNS or the metabolic state manipulation, we fit reinforcement learning models to participants’ choice behavior as previously detailed (Guitart-Masip et al., 2014, 2012b; Kühnel et al., 2020) using hierarchical expectation maximization (Huys et al., 2011). To avoid artificially inducing group differences, we assumed one group-level distribution for each parameter and estimated parameters for each session as independent data. Briefly, we estimated a five-parameter model including one learning rate, a reward sensitivity, the noisiness of choices, a Pavlovian choice bias, and a go bias, the same model as reported previously (Kühnel et al., 2020).

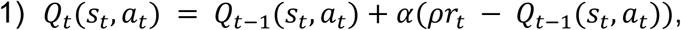

The model assumes that participants learn stimulus (s) specific action (a) values (Q) that are updated according to the Rescorla-Wagner rule in each trial (t). The speed of learning is determined by the learning rate (α ɛ [0,1]) and the importance of rewards (coded as-1, 0, 1) is scaled by the reward sensitivity (ρ ɛ [0,Inf]). Additionally, for each stimulus, action-independent values (V) are learned using the same rule to indicate whether a stimulus is associated with rewards or punishments.

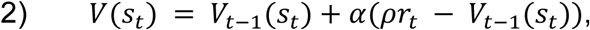

Then, action weights for the stimulus in the current trial are computed combining the learned action values (Q) with the stimulus values (V) weighted by the Pavlovian choice bias (π ɛ [0,Inf]) that increases the go probability in win trials (vs. punishment trials) and a constant go bias (b ɛ [-Inf,Inf]):

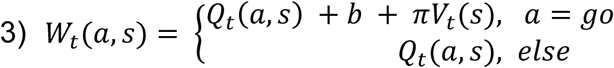

Last, the action probabilities for each trial are estimated using a softmax function with the action weights (*W*) and an additional noise parameter (lapse, ξ ∈ [0, 1]).

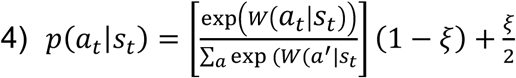

To account for non-normal distributions of parameters from the computational model, differences in parameter estimates between the tVNS and sham conditions were tested using bootstrapping (10,000 resamples). For this analysis, we only used complete datasets. For the associations of model parameters with hunger and satiety, we used the same linear mixed-effects model using the parameters of the computational model as outcome. To compare the results of the current study with our previous results (Kühnel et al., 2020), we used independent sample t-tests on tVNS-induced differences in learning rates (tVNS - sham) comparing the previous data with either the data from the sessions with a milkshake load or from the water condition.

### Statistical threshold and software

For all main outcomes, we used a significance threshold of *p* <.05 (two-tailed). Data analyses have been performed using MATLAB v2022a and R v4.4.2 (Posit team, 2025; R Core Team, 2024). For statistical modeling, we used the lmer function of the lmerTest package in R (Kuznetsova et al., 2017), which estimates degrees of freedom using the Satterthwaite approximation. For data visualization, we used ggplot2 (Wickham, 2016), ggridges (Wilke, 2024a), cowplot (Wilke, 2024b), tidybayes (Kay, 2024), and viridis (Garnier et al., 2024).

### Results tVNS reduces Pavlovian bias after a caloric load

First, we analyzed the typical task-related effects on choice accuracy. In line with previous findings, participants performed better in go versus no-go trials (“go bias”), *t*(53.49) = 4.98*, p* <.001 (Fig. 2A), and in trials with congruent action-outcome contingencies (i.e., go in win and no-go in punishment trials) versus incongruent trials (“Pavlovian bias”), Win × Go: *t*(53.18) = 7.87, *p* <.001 (Fig. 2C). In addition, we also observed an effect of reward valence on choice accuracy: participants performed worse in win trials compared to punishment trials, *t*(53.38) =-3.22, *p* =.002 (Fig. 2B). Participants successfully learnt contingencies, indicated by improved performance over trials, *t*(52.86) = 10.01, *p* <.001 (Fig. 2D). Moreover, performance increased over sessions, *t*(105.10) = 10.50, *p* <.001. To assess whether Pavlovian biases improve or reduce overall performance, we correlated random effect estimates of the Go × Win interaction with the average percentage of correct choices. The resulting negative association (*r* =-.36, *p* =.007) suggests that better overall performance is associated with a weaker Pavlovian bias.

**Figure 2:**
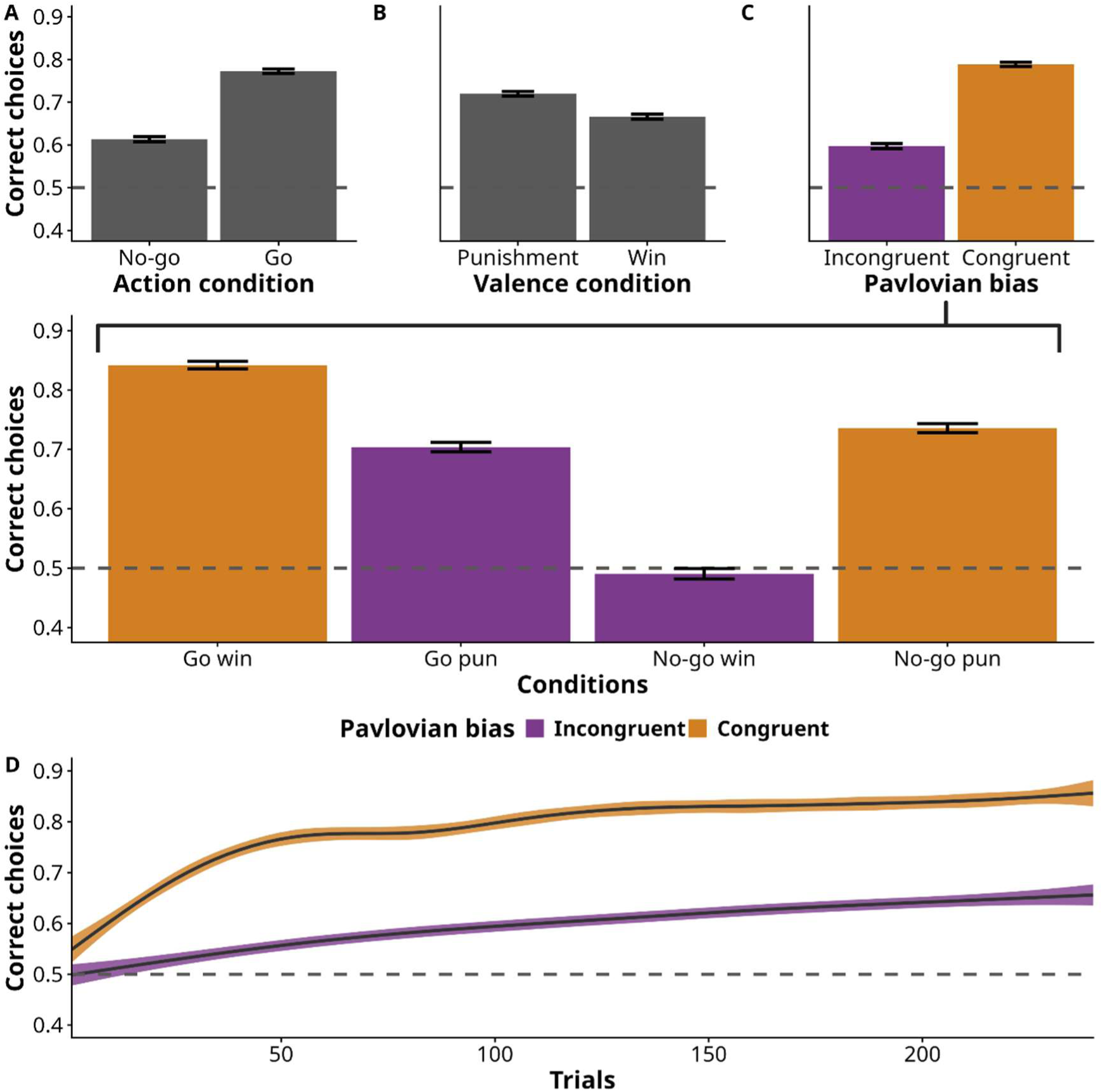
Participants demonstrate Pavlovian bias and improvements over trials. A: Participants showed improved performance in go trials compared to no-go trials, *t*(53.49) *=* 4.98*, p* <.001. B: Participants performed worse in win compared to punishment trials, *t*(53.38) =-3.22, *p* =.002. C: Participants showed improved performance in trials with congruent action-outcome contingencies (i.e., go in win and no-go in punishment trials) demonstrating Pavlovian bias, Win × Go: *t*(53.18) = 7.87, *p* <.001. D: Participants’ performance improved over trials, particularly in congruent trials, *t*(52.86) = 10.01, *p* <.001.

Next, we investigated the effects of tVNS and caloric load on choice accuracy. Although there were no main effects of stimulation, *t*(51.19) *=*-0.66*, p* =.51), or caloric load, *t*(53.32) = 0.09, *p* =.93 (Fig. 3A), tVNS modulated choice accuracy in go trials in a state-dependent manner (Go × Stimulation × Load: *t*(48.67) =-2.10, *p* =.041), leading to improved performance after water vs. milkshake intake. Likewise, tVNS modulated the Go × Win interaction by improving performance in congruent trials (e.g., go to win) after water intake while reducing performance in congruent trials after milkshake intake (expanded term with Stimulation x Load: *t*(53.39) =-2.12, *p* =.038, Fig. 3C).

**Figure 3:**
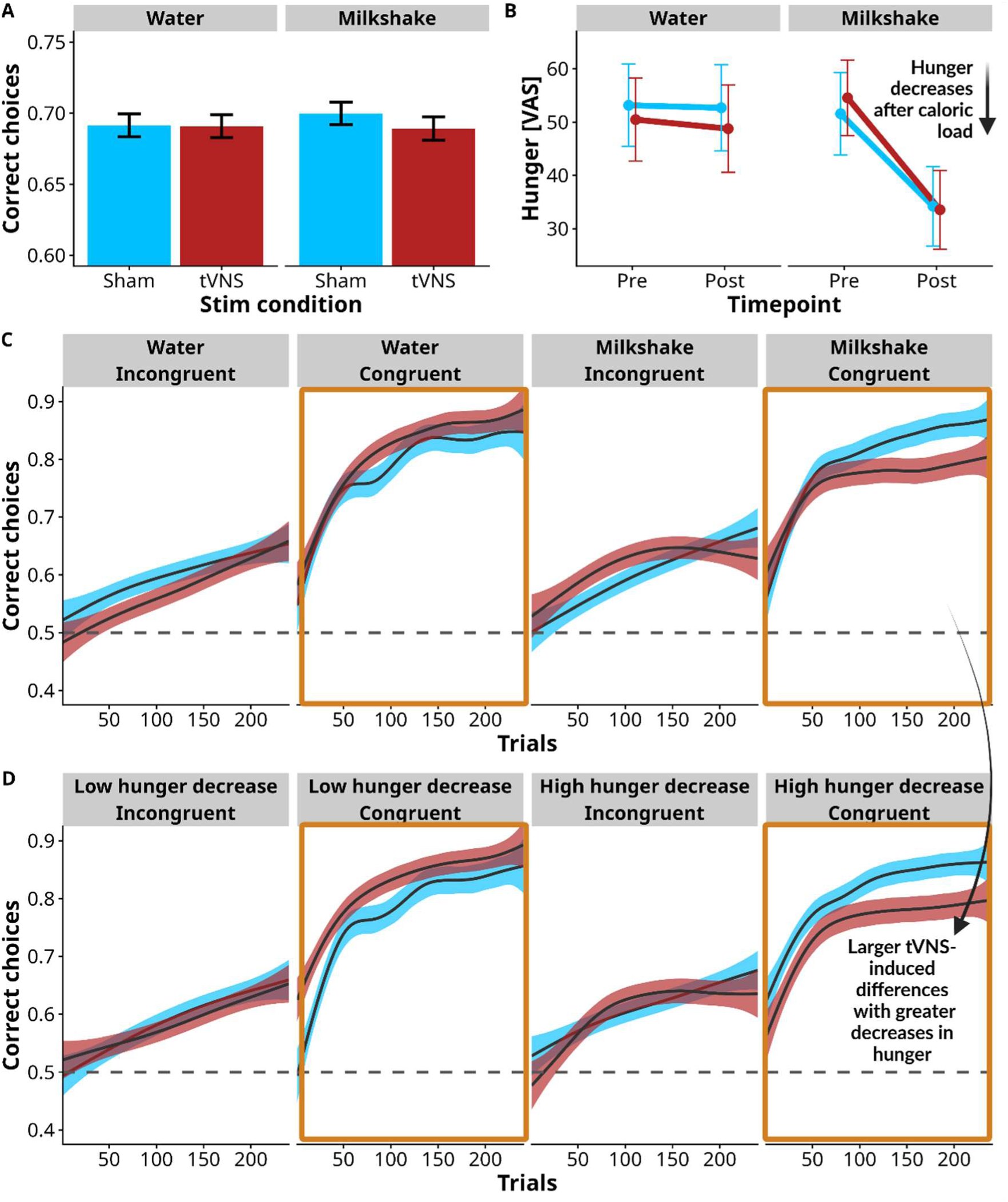
tVNS modulates Pavlovian bias in a state-dependent manner. A: Choice accuracy was not affected by tVNS (*t*(51.19) =-0.66*, p* =.51) or the caloric load (*t*(53.32) = 0.09*, p* =.93). B: ΔHunger decreased more pronounced after the milkshake vs water (*t*(83.17) =-6.41, *p* <.001). C: tVNS modulated the Go × Win interaction by improving performance in congruent trials (e.g., go to win) after water intake and reducing performance in congruent trials after milkshake intake (expanded term with Stimulation × Load: *t*(53.39) =-2.12, *p* =.038). D: Using ΔHunger we observed tVNS-induced improvements in performance in congruent trials (i.e., increased Pavlovian bias, Go × Win) and decreases in performance in incongruent trials (e.g., no-go to win) if participants were hungrier (i.e., after the water load), Stimulation × ΔHunger: *t*(162.18) = 2.63, *p* =.009).

### Metabolic state drives tVNS-induced changes in Pavlovian bias

To investigate whether tVNS-induced effects on Pavlovian bias were driven by interoception of metabolic states, we used changes in hunger and satiety ratings (ΔHunger; ΔSatiety, post load – pre load) instead of the load condition (milkshake vs. water). As expected, milkshake intake led to a greater reduction in hunger (i.e., negative ΔHunger) compared to water, *t*(83.17) =-6.41, *p* <.001 (Fig. 3B) and an increase in satiety (i.e., ΔSatiety), *t*(53.13) = 6.35, *p* <.001. Using ΔHunger, we observed tVNS-induced improvements in performance in congruent trials (i.e., increased Pavlovian bias, Go × Win) and decreases in performance in incongruent trials (e.g., no-go to win) if participants were hungrier after the load, Stimulation × ΔHunger: *t*(162.18) = 2.63, *p* =.009 (Fig. 3D). However, using ΔSatiety did not modulate the effect of tVNS, Stimulation × ΔSatiety: *t*(161.22) =-1.72, *p* =.087. Model comparisons using AIC and BIC indicated that individual changes in hunger ratings best accounted for the observed effects (ΔBIC = - 4.49, SI2).

### Computational modeling of Pavlovian bias

To map state-dependent changes in choice behavior onto distinct computational parameters, we used the same 5-parameter model (learning rate, reward sensitivity, Pavlovian bias, go bias, and noise) that explained the data well in previous work (Kühnel et al., 2020). To capture the load-dependent effect of the stimulation, we bootstrapped the differences (ΔPavBias = (tVNS_load_ – sham_load_) – (tVNS_water_ – sham_water_)) reflecting the interaction of load condition and stimulation. First, we compared the effects of tVNS on the learning rate in the water condition (i.e., fasting state) with the reduction in learning in our previous study (Kühnel et al., 2020). Mirroring earlier results, the learning rate with tVNS vs. sham was nominally lower in the water condition, ΔLearningRate =-0.041, *p*_boot_ =.069, and the observed effect did not differ significantly from the previous result, *t*(79) = 1.2, *p* =.25 (Fig. 4B), in contrast to the tVNS-induced change in the milkshake condition, *t*(79) = 2.49, *p* =.015. Second, in line with the model-agnostic results, tVNS induced increases in the PavBias parameter in the water condition, but led to decreases in the milkshake condition. However, the effect did not reach significance, Δlog(PavBias) =-0.33, *p*_boot_ =.065 (Fig. 4A,C), although the PavBias correlated strongly with the random effects estimates of the Go × Win interaction term from the mixed-effects model (*r* =.74, *p* <.001). To investigate whether effects were stronger when using hunger ratings, we used a comparable linear mixed-effects model as for the hunger effects for the Go × Win slope. Mirroring the results comparing the load conditions, tVNS effects did not significantly depend on changes in hunger ratings, *t*(144) = 1.68, *p* =.095, or satiety ratings, *t*(135) =-1.26, *p* =.21. No other parameter showed a (state-dependent) tVNS effect (*p*s >.19, Fig. 4A).

**Figure 4:**
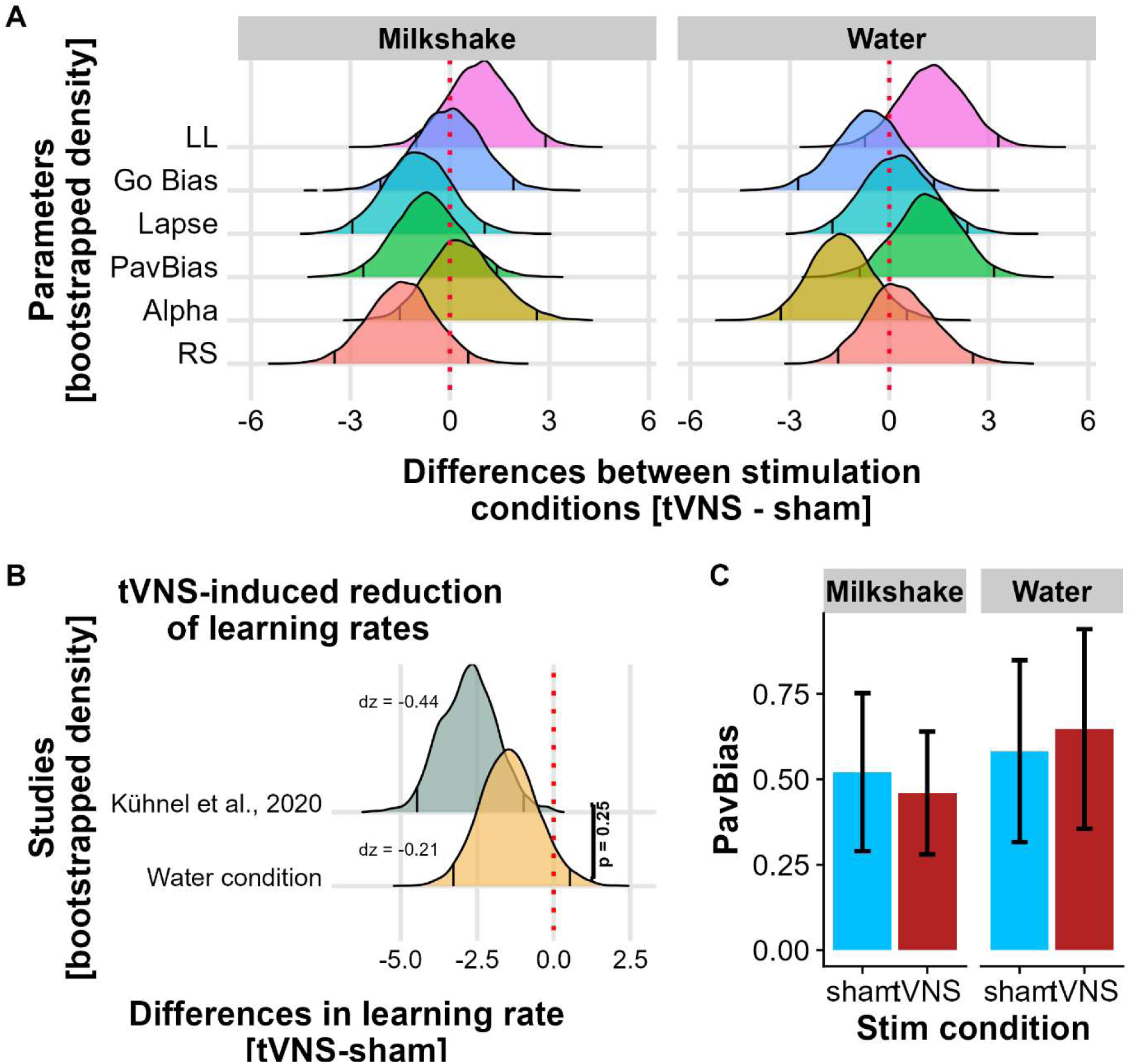
**Computational modeling of learning behavior**. A: Bootstrapped (10,000 resamples) differences (tVNS - sham) of tVNS-induced changes in learning parameters show that Pavlovian bias (PavBias) decreased during tVNS after the milkshake load, whereas it increased after the water load (Δlog(PavBias) =-0.33, *p*_boot_ =.065). B: We compared the effects of tVNS on the learning rate in the fasting condition (i.e., after the water load) with the reduction in learning in our previous study (Kühnel et al., 2020): The learning rate for tVNS vs. sham was nominally lower in the water condition (ΔLearningRate =-0.041, *p*_boot_ =.069) and the observed effect did not differ significantly from the previous result, *t*(79) = 1.2, *p* =.25). C: Effect of tVNS and metabolic conditions on modeled Pavlovian bias (PavBias), error bars show 95% confidence intervals.

### Sensitivity and control analyses

To assess successful blinding, participants were asked after each session, “What stimulation condition do you think you had today?”. Participants guessed the correct condition with chance probability (205 guesses: 112 correct guesses, accuracy 54.63 %, *p*_binom_ =.10), demonstrating successful blinding. In addition, correct guesses did not increase over sessions, *b* = 0.11, *SE* = 0.17, *z* = 0.62, *p* =.54. To account for demand effects (i.e., expectation of treatment effect), we included a regressor for participants’ guessed stimulation condition in the load-condition model (effect-coded:−0.5 = sham, 0.5 = tVNS), with random intercepts and random slopes for each participant (for model equations, see SI3). In this model, the effect of the stimulation and load condition on the Go × Win interaction became even more pronounced compared to the model without this regressor, *t*(53.27) = −2.70, *p* =.009. In contrast, tVNS guesses were associated with a lower PavBias in the computational model, *t*(140.39) =-2.29, *p* =.024. Intriguingly, once we accounted for condition-related guesses, the difference between tVNS water and milkshake load conditions became significant (*p* =.030), suggesting that tVNS enhanced the caloric load effect compared to sham (*p* =.54). As tVNS-induced effects on reinforcement learning have previously been described to partly depend on BMI (Kühnel et al., 2020), we tested the interaction with BMI in our current sample, but did not find any significant effect, Stimulation × BMI: *t*(50.98) =-1.11, *p* =.27.

## Discussion

Recent evidence indicates that metabolic states gate the effects of vagal afferent stimulation and may thereby modulate tVNS-induced effects on reward-related behavior (Teckentrup and Kroemer, 2024). Consequently, we investigated whether tVNS-induced effects on performance in a valenced go/no-go reinforcement learning task depend on metabolic state using a milkshake vs. water preload. In line with the hypothesized state-dependent effects, tVNS amplified the Pavlovian bias in a fasting state while it reduced the Pavlovian bias when hunger decreased after the load. We conclude that tVNS exerts state-dependent effects on motivational biases that shape reinforcement learning, adding to the growing evidence that bodily states should be considered to optimize the application of tVNS.

Although VNS is typically administered independent of internal or external states, we show that tVNS amplified Pavlovian bias in a fasted state, and reduced Pavlovian bias after energy intake. A Pavlovian choice bias (i.e., approaching rewards and avoiding punishments (Hershberger, 1986)) may be adaptive in a state when energy resources are scarce. Consequently, rodents demonstrate higher approach behavior towards food and faster acquisition of a cue-food conditioned response when fasted compared to when fed (Anversa et al., 2025; Lockie et al., 2017; Shteyn et al., 2025). Intriguingly, the tVNS-induced modulatory effect was best captured by individual changes in hunger ratings but not satiety ratings. Distinct neural pathways have been demonstrated for food seeking and suppression in rodents (Chen et al., 2020; Shen et al., 2025), suggesting that our results may have been driven primarily by gating the motivating effects of hunger that may facilitate valence-contingent actions. To conclude, state-dependent effects of tVNS on motivational biases in learning support the idea that VNS modulates behavior by gating internal signals of demand (Teckentrup and Kroemer, 2024).

Prior studies investigating tVNS-induced effects on reinforcement learning have predominantly used left or bilateral tVNS (Çakır et al., 2025; D’Agostini et al., 2021; Kühnel et al., 2020; Weber et al., 2021). In addition to lateralized effects, fasting has been shown to enhance dopamine tone (Ostlund et al., 2011). In line with previous work (Kühnel et al., 2020; Thanarajah et al., 2025), right tVNS during a fasting state (water condition) reduced the learning rate. This observation is compatible with an increased dopamine tone, leading to a reduced signal-to-noise ratio of phasic prediction error signals (Beeler, 2012; Hamid et al., 2016; Kroemer et al., 2019). Elevated dopamine tone has also been associated with increased motivation in general (Beierholm et al., 2013; Hamid et al., 2016; Salamone et al., 2016). In line with this association and with previous pharmacological interventional studies (Guitart-Masip et al., 2012a), we observed improved performance on go trials with right tVNS after water intake. However, state-dependent effects of tVNS on Pavlovian bias might be best explained by the gating of internal signals (Teckentrup and Kroemer, 2024) rather than changes in dopamine signaling. In addition to dopamine, modulation of noradrenaline by VNS (Collins et al., 2021; Giraudier et al., 2023; Mridha et al., 2021) may also play a role in regulating vigor (Varazzani et al., 2015), learning (O’Callaghan, 2025), and motivational biases (Campese et al., 2017; Pasquariello et al., 2018). For example, a recent study has shown that pupil dilation (a proxy of phasic noradrenergic signaling modulated by tVNS; Pervaz et al., 2025) reflects effortful action invigoration as needed to overcome inhibition (Algermissen and den Ouden, 2024). Consequently, our results may suggest that tVNS induces behavioral effects via changes in monoaminergic signaling, but such changes are dependent on a motivational state of need (fasting/hunger).

Despite the strength of systematically varying metabolic state during right-sided tVNS in a within-subject design, our study also has several limitations. First, although prior research has linked VNS and dopamine signaling (Brougher et al., 2021; Han et al., 2018), vagal afferents induce non-selective neuromodulatory effects (Collins et al., 2021; Hulsey et al., 2016; Roosevelt et al., 2006). Since direct assessments of neurotransmitter release are limited in humans, specifically in the midbrain or brainstem, further studies investigating tVNS-induced changes in neurotransmission are necessary to understand the likely mechanisms. Second, while hunger ratings provide a proxy of the individuals’ interoception of the metabolic effects of the caloric load (Kaduk et al., 2025; Lemmens et al., 2011; Schultes et al., 2003), future studies may incorporate physiological measures (e.g., changes in glucose or insulin levels). Third, we applied conventional tVNS throughout the task, independent of specific external events (e.g., reward outcomes). In contrast, VNS-induced improvements in motor learning or fear extinction are dependent on VNS being paired with specific events (Bowles et al., 2022; Souza et al., 2022). Using pulsed tVNS may modulate phasic monoaminergic signals (Pervaz et al., 2025) instead of altering monoaminergic tone. Therefore, an outcome-or action-paired pulsed tNVS protocol might be more effective to induce learning improvements. Fourth, although we observed significant differences in the model-agnostic analyses, the computational model only showed a trend. One reason for this discrepancy might be differences in how the hierarchy of the data was modeled since the computational model did not consider the nesting within participants, likely leading to a loss of power. However, expanding the hierarchical structure of the computational model may require more repetitions within participants compared to mixed-effects models of choices.

To conclude, we demonstrated a state-dependent effect of right tVNS on Pavlovian choice bias, best captured by individual load-induced changes in hunger but not satiety ratings. These findings contribute to a growing body of evidence supporting a bottom-up influence of vagal afferents on motivational approach behavior and reinforcement learning, tuning goal-directed behavior according to metabolic state (Teckentrup and Kroemer, 2024). Our results further emphasize the critical role of metabolic state in shaping tVNS-induced effects, which may guide future research aimed at optimizing the application of tVNS in both experimental and clinical contexts. Based on our work and earlier findings (Teckentrup and Kroemer, 2024), it seems imperative to consider appropriate metabolic states for the administration of tVNS to elicit intended neuromodulatory effects that lead to changes in motivation and learning.

## Acknowledgement

We thank Ina Faupel, Lina Knees, Charlotte Landmann, and Anne Schiller for help with data acquisition. The study was supported by the Daimler and Benz Foundation postdoctoral scholarship 32-04/19 awarded to NBK. In addition, it was supported by DFG 493623632, DFG KR 4555/7-1, KR 4555/9-1, and KR 4555/10-1, BONFOR O-128.0101.

## Author contributions

NBK was responsible for the study concept and design. YW, NG, & CS collected data under supervision by NBK. AK & NBK conceived the method, and AK & YW processed the data. AK and LK performed the data analysis, and CS and NBK contributed to the analyses and data visualizations. LK, AK, CS, & NBK wrote the manuscript. All authors contributed to the interpretation of findings, provided critical revision of the manuscript for important intellectual content and approved the final version for publication.

