## supplements for "Non-invasive vagus nerve stimulation modulates Pavlovian bias in a state-dependent manner"

**Corresponding author***

### SI1. Model-agnostic output

**Table SI1**

| | b | SE | df | t | p |
|---|---|---|---|---|---|
| **(Intercept)** | 0.70 | 0.02 | 52.83 | 34.96 | < 0.001 |
| **cStimulation** | -0.01 | 0.01 | 51.19 | -0.66 | .51 |
| **cWin** | -0.05 | 0.02 | 53.38 | -3.22 | .002 |
| **cGo** | 0.15 | 0.03 | 53.49 | 4.98 | < 0.001 |
| **cLoad** | 0.001 | 0.02 | 53.32 | 0.09 | .93 |
| **cSession** | 0.09 | 0.01 | 105.10 | 10.50 | < 0.001 |
| **cTrial** | 0.06 | 0.01 | 52.86 | 10.01 | < 0.001 |
| **cStimulation × cWin** | -0.01 | 0.02 | 50.92 | -0.31 | .76 |
| **cStimulation × cGo** | -0.01 | 0.04 | 52.95 | -0.24 | .81 |
| **cWin × cGo** | 0.38 | 0.05 | 53.18 | 7.87 | < 0.001 |
| **cStimulation × cLoad** | -0.005 | 0.02 | 48.83 | -0.22 | .83 |
| **cWin × cLoad** | 0.02 | 0.02 | 48.97 | 1.52 | .14 |
| **cGo × cLoad** | -0.03 | 0.03 | 53.39 | -0.77 | .44 |
| **cStimulation × cWin × cGo** | -0.01 | 0.06 | 50.74 | -0.16 | .88 |
| **cStimulation × cWin × cLoad** | 0.07 | 0.03 | 47.69 | 1.95 | .058 |
| **cStimulation × cGo × cLoad** | -0.15 | 0.07 | 48.67 | -2.10 | .041 |
| **cWin × cGo × cLoad** | -0.08 | 0.05 | 52.10 | -1.58 | .12 |
| **cStimulation × cWin × cGo × cLoad** | -0.21 | 0.10 | 53.39 | -2.12 | .038 |

Model equation: lmer(Correct ~ cStimulation × cWin × cGo × cLoad + cSession + cTrial + (1 + cStimulation × cWin × cGo × cLoad + cTrial | ID) data)

**Table SI2**

| | b | SE | df | t | p |
|---|---|---|---|---|---|
| **(Intercept)** | 0.70 | 0.02 | 52.77 | 38.42 | <0.001 |
| **cStimulation** | -0.01 | 0.01 | 50.59 | -0.61 | .54 |
| **cWin** | -0.05 | 0.02 | 53.44 | -3.22 | .002 |
| **cGo** | 0.15 | 0.03 | 53.68 | 4.98 | <0.001 |
| **cLoad** | 0.002 | 0.01 | 53.50 | 0.14 | .89 |
| **cSession** | 0.09 | 0.01 | 105.00 | 10.72 | <0.001 |
| **cTrial** | 0.06 | 0.01 | 52.99 | 10.02 | <0.001 |
| **cSex** | -0.04 | 0.02 | 49.50 | -2.24 | .030 |
| **cBMI** | -0.003 | 0.002 | 49.13 | -1.67 | .10 |
| **cAge** | -0.003 | 0.001 | 49.03 | -4.40 | <0.001 |
| **cStimulation × cWin** | -0.01 | 0.02 | 50.72 | -0.31 | .76 |
| **cStimulation × cGo** | -0.01 | 0.04 | 52.83 | -0.24 | .81 |
| **cWin × cGo** | 0.38 | 0.05 | 53.38 | 7.87 | <0.001 |
| **cStimulation × cLoad** | -0.01 | 0.02 | 49.42 | -0.23 | .82 |
| **cWin × cLoad** | 0.02 | 0.02 | 49.22 | 1.54 | .13 |
| **cGo × cLoad** | -0.03 | 0.03 | 53.28 | -0.77 | .44 |
| **cStimulation × cWin × cGo** | -0.01 | 0.06 | 50.16 | -0.21 | .84 |
| **cStimulation × cWin × cLoad** | 0.07 | 0.03 | 47.79 | 1.97 | .055 |
| **cStimulation × cGo × cLoad** | -0.15 | 0.07 | 48.86 | -2.12 | .039 |
| **cWin × cGo × cLoad** | -0.08 | 0.05 | 52.44 | -1.66 | .10 |
| **cStimulation × cWin × cGo × cLoad** | -0.21 | 0.10 | 53.66 | -2.17 | .035 |

Model equation: lmer (Correct ~ cStimulation × cWin × cGo × cLoad + cSession + cTrial + cSex + cBMI + cAge + (1 + cStimulation × cWin × cGo × cLoad + cTrial | ID) data)

**Table SI3**

| | b | SE | df | t | p |
|---|---|---|---|---|---|
| **(Intercept)** | 0.70 | 0.02 | 52.65 | 35.99 | <0.001 |
| **cStimulation** | -0.01 | 0.01 | 50.54 | -0.69 | .49 |
| **cWin** | -0.05 | 0.02 | 52.52 | -3.24 | .002 |
| **cGo** | 0.15 | 0.03 | 52.62 | 4.95 | <0.001 |
| **cLoad** | 0.0007 | 0.01 | 52.12 | 0.05 | .96 |
| **cBMI** | -0.01 | 0.002 | 51.05 | -2.73 | .009 |
| **cSession** | 0.09 | 0.01 | 103.30 | 10.23 | <0.001 |
| **cTrial** | 0.06 | 0.01 | 52.90 | 10.01 | <0.001 |
| **cStimulation × cWin** | -0.01 | 0.02 | 50.03 | -0.31 | .76 |
| **cStimulation × cGo** | -0.01 | 0.04 | 52.08 | -0.21 | .83 |
| **cWin × cGo** | 0.38 | 0.05 | 52.62 | 7.91 | <0.001 |
| **cStimulation × cLoad** | -0.003 | 0.02 | 47.70 | -0.16 | .87 |
| **cWin × cLoad** | 0.02 | 0.02 | 47.69 | 1.51 | .14 |
| **cGo × cLoad** | -0.03 | 0.03 | 52.17 | -0.74 | .46 |
| **cStimulation × cBMI** | -0.002 | 0.002 | 50.98 | -1.11 | .27 |
| **cWin × cBMI** | 0.004 | 0.003 | 52.20 | 1.50 | .14 |
| **cGo × cBMI** | 0.004 | 0.01 | 52.67 | 0.70 | .49 |
| **cLoad × cBMI** | -0.004 | 0.002 | 51.66 | -1.59 | .12 |
| **cStimulation × cWin × cGo** | -0.003 | 0.05 | 50.57 | -0.05 | .96 |
| **cStimulation × cWin × cLoad** | 0.07 | 0.03 | 45.64 | 1.96 | .056 |
| **cStimulation × cGo × cLoad** | -0.15 | 0.07 | 47.55 | -2.12 | .039 |
| **cWin × cGo × cLoad** | -0.07 | 0.05 | 50.51 | -1.44 | .16 |
| **cStimulation × cWin × cBMI** | -0.001 | 0.003 | 49.66 | -0.33 | .75 |
| **cStimulation × cGo × cBMI** | 0.01 | 0.01 | 52.00 | 0.89 | .38 |
| **cWin × cGo × cBMI** | -0.01 | 0.01 | 52.22 | -1.82 | .075 |
| **cStimulation × cLoad × cBMI** | 0.004 | 0.003 | 48.71 | 1.14 | .26 |
| **cWin × cLoad × cBMI** | -0.0007 | 0.003 | 47.12 | -0.28 | .78 |
| **cGo × cLoad × cBMI** | 0.004 | 0.01 | 52.23 | 0.65 | .52 |
| **cStimulation × cWin × cGo × cLoad** | -0.22 | 0.10 | 51.65 | -2.26 | .028 |
| **cStimulation × cWin × cGo × cBMI** | 0.02 | 0.01 | 49.98 | 1.74 | .088 |
| **cStimulation × cWin × cLoad × cBMI** | -0.01 | 0.01 | 45.93 | -1.17 | .25 |
| **cStimulation × cGo × cLoad × cBMI** | -0.01 | 0.01 | 47.19 | -0.72 | .48 |
| **cWin × cGo × cLoad × cBMI** | 0.01 | 0.01 | 50.54 | 0.82 | .42 |
| **cStimulation × cWin × cGo × cLoad × cBMI** | -0.03 | 0.02 | 51.11 | -1.70 | .095 |

Model equation: lmer(Correct ~ cStimulation × cWin × cGo × cLoad × cBMI + cSession + cTrial + (1 + cStimulation × cWin × cGo × cLoad + cTrial |ID), data)

**Table SI4**

| | b | SE | df | t | p |
|---|---|---|---|---|---|
| **(Intercept)** | 0.39 | 0.05 | 50.54 | 8.39 | < 0.001 |
| **cStimulation** | -0.01 | 0.05 | 47.40 | -0.26 | .80 |
| **cΔHunger** | 0.03 | 0.13 | 32.95 | 0.20 | .84 |
| **cStimulation × cΔHunger** | 0.58 | 0.22 | 162.18 | 2.63 | .009 |

Model equation: lmer(cWin × cGo ~ cStimulation × cΔHunger  (1 + cStimulation + cΔHunger |ID), data)

**Table SI5**

| | b | SE | df | t | p |
|---|---|---|---|---|---|
| **(Intercept)** | 0.38 | 0.05 | 53.39 | 8.16 | < 0.001 |
| **cStimulation** | -0.005 | 0.05 | 59.46 | -0.10 | .92 |
| **cΔSatiety** | -0.04 | 0.13 | 65.99 | -0.32 | .75 |
| **cStimulation × cΔSatiety** | -0.38 | 0.22 | 161.22 | -1.72 | .087 |

Model equation: lmer(cWin × cGo ~ cStimulation × cΔSatiety  (1 + cStimulation + cΔSatiety |ID), data)

**SI2. Model Comparison**

**Table SI6**

| model | df | BIC |
|---|---|---|
| **cLoad** | 11 | 296.88 |
| **cΔhunger** | 11 | 292.39 |
| **cΔSatiety** | 11 | 295.41 |
| | | **AIC** |
| **cLoad** | 11 | 260.06 |
| **cΔHunger** | 11 | 255.57 |
| **cΔSatiety** | 11 | 258.60 |

**SI3. Sensitivity and control analysis**

**Table SI7**

| | b | SE | df | t | p |
|---|---|---|---|---|---|
| **(Intercept)** | 0.70 | 0.02 | 50.61 | 33.44 | < 0.001 |
| **cStimulation** | -0.02 | 0.01 | 32.58 | -1.54 | .13 |
| **cWin** | -0.05 | 0.02 | 52.80 | -3.08 | .003 |
| **cGo** | 0.15 | 0.03 | 53.56 | 4.70 | < 0.001 |
| **cLoad** | 0.01 | 0.02 | 50.97 | 0.47 | .64 |
| **cGuesses** | 0.005 | 0.01 | 24.14 | 0.31 | .76 |
| **cSession** | 0.09 | 0.01 | 71.85 | 12.32 | < 0.001 |
| **cTrial** | 0.06 | 0.01 | 53.08 | 9.82 | < 0.001 |
| **cStimulation × cWin** | -0.01 | 0.02 | 48.80 | -0.58 | .57 |
| **cStimulation × cGo** | -0.02 | 0.04 | 52.79 | -0.46 | .65 |
| **cWin × cGo** | 0.37 | 0.05 | 54.08 | 7.45 | < 0.001 |
| **cStimulation × cLoad** | 0.02 | 0.03 | 39.56 | 0.91 | .37 |
| **cWin × cLoad** | 0.02 | 0.01 | 46.31 | 1.14 | .26 |
| **cGo × cLoad** | -0.03 | 0.03 | 54.13 | -0.97 | .34 |
| **cStimulation × cWin × cGo** | -0.01 | 0.05 | 49.45 | -0.13 | .90 |
| **cStimulation × cWin × cLoad** | 0.07 | 0.03 | 43.07 | 1.93 | .061 |
| **cStimulation × cGo × cLoad** | -0.17 | 0.07 | 47.94 | -2.41 | .020 |
| **cWin × cGo × cLoad** | -0.08 | 0.05 | 49.11 | -1.64 | .11 |
| **cStimulation × cWin × cGo × cLoad** | -0.26 | 0.10 | 53.27 | -2.70 | .009 |

Model equation: lmer(Correct ~ cStimulation × cWin × cGo × cLoad + cGuesses + cSession + cTrial + (1 + cStimulation × cWin × cGo × cLoad + cGuesses + cTrial |ID), data)

**Table SI8**

| | b | SE | df | t | p |
|---|---|---|---|---|---|
| **(Intercept)** | 0.55 | 0.09 | 53.43 | 5.86 | < 0.001 |
| **cStimulation** | 0.009 | 0.11 | 54.93 | 0.09 | .93 |
| **cLoad** | -0.13 | 0.09 | 58.86 | -1.44 | .15 |
| **cGuesses** | -0.19 | 0.08 | 140.39 | -2.29 | .024 |
| **cSession** | 0.10 | 0.07 | 119.22 | 1.43 | .16 |
| **cStimulation × cLoad** | -0.21 | 0.12 | 97.73 | -1.73 | .088 |

Model equation: lmer(PavBias ~ cStimulation × cLoad + cGuesses + cSession + (1 + cStimulation + cLoad |ID), data)
